# High-density exploration of activity states in a multi-area brain model

**DOI:** 10.1101/2023.05.18.541285

**Authors:** David Aquilué-Llorens, Jennifer S. Goldman, Alain Destexhe

**Affiliations:** Paris-Saclay University, CNRS, Paris-Saclay Institute of Neuroscience (NeuroPSI), 91400 Saclay, France; Starlab Barcelona SL, Neuroscience BU, Av Tibidabo 47 bis, Barcelona, Spain

**Keywords:** Cerebral cortex, Computational models, Asynchronous states, Information processing, Whole-brain model, Parameter exploration, Brain states, High-performance computing

## Abstract

Biophysically-grounded whole-brain models were built recently using tractography data to interconnect multiple mesoscopic models, which can simulate the dynamics of neuronal populations with only a few equations. Mean-field models of neural populations, specifically the Adapting AdEx mean-field, was used for this purpose because it can integrate key biophysical mechanisms such as spike-frequency adaptation and its regulation at cellular scales, to the emergence of brain-scale dynamics. Using this approach, with the Virtual Brain (TVB) environment, it has been possible to model the macroscopic transitions between brain states, described by variation in brain-scale dynamics between asynchronous and rapid dynamics during conscious brain states, and synchronized slow-waves, with Up-and-Down state dynamics during unconscious brain states, emerging from mechanisms at the cellular level. Transitions between brain states are driven by changes in neuromodulation that can be due to intrinsic regulation during sleep-wake cycles or extrinsic factors such as anesthetics, which, in turn, affect spike-frequency adaptation. Here, we perform a dense grid parameter exploration of the TVB-AdEx model, making use of High Performance Computing, to thoroughly explore the properties of this model. We find that there is a remarkable robustness of the effect of adaptation to induce synchronized slow-wave activity. Moreover, the occurrence of slow waves is often paralleled with a closer relation between functional and structural connectivity. We find that hyperpolarization can also generate unconscious-like synchronized Up and Down states, which may be a mechanism underlying the action of anesthetics. We conclude that the parameter space of the TVB-AdEx model reveals features identified experimentally in sleep and anesthesia.

## 1. Introduction

Consciousness is a fundamental aspect of human existence, it is experiencing, having a thought, being aware of feeling an emotion. Unconsciousness is not unfamiliar either: we naturally lose consciousness when going into dreamless (NREM) sleep or medically through external factors like general anesthesia for surgery or brain injury. However, defining what consciousness is exactly, from what processes it emerges, and even the possibility of measuring it, is still an open and extremely complex philosophical and scientific question. Furthermore, disorders of consciousness, such as coma or unresponsive wakefulness syndrome, as well as disorders of the wake-sleep cycle, such as insomnia, take a great toll on society [1, 2] and approximately 200 million surgical procedures are carried out under anesthesia each year [3]. Thus, the study of consciousness is also relevant from the medical point of view.

Although there does not seem to be particular brain regions uniquely responsible for consciousness [4], it is possible to roughly categorize the macroscopic, global activity of the brain into conscious (during dreaming or being awake) and unconscious (during deep sleep and under general anesthesia) states [5].

On the one hand, noninvasive, macroscopic recordings of conscious states are typically characterized by low amplitude, complex and disorganized signals over the brain, that function at relatively high frequencies. On the other hand, unconscious states exhibit higher amplitude but simpler, synchronous, low frequency oscillating activity [5, 6].

Additionally, interactions between brain areas are strongly mediated by the anatomical white matter tracts during unconscious states, while, in conscious states, more complex brain activity explores a richer repertoire of functional configurations that are less constrained by brain structure [7, 8]. Finally, the complex and disorganized activity of conscious states allows the brain to be more responsive to external stimuli, unlike unconscious states, during which responses to stimuli do not propagate much throughout the cortex [9].

There are also observable differences between conscious and unconscious states at the scale of single neurons. During consciousness, neurons have been seen to exhibit sustained but sparse and irregular firing patterns, termed Asynchronous Irregular (AI). In contrast, during periods of unconsciousness, neurons oscillate between hyperpolarized (Down) states and depolarized and AI-like firing (Up) states [5, 6, 10].

Differences in neuron behavior between brain states can result from changes in neuromodulator concentration, which affect spike-frequency adaptation. Spike-frequency adaptation is a self-inhibiting process in which the neuron experiences a decrease in its firing frequency the more it fires, sometimes to the point of silencing the neuron [11]. During conscious states, concentrations of neuromodulators such as acetylcholine are higher [12], which suppresses spike frequency adaptation and facilitates AI firing patterns. During unconscious states, such neuromodulator concentrations are lower, increasing spike frequency adaptation, forcing neurons into alternating periods of silence and activity.

Understanding how the macroscopic global dynamics of conscious and unconscious brain states relate to the microscopic neuromodulatory processes that take place at the scale of single neurons is an important problem that can help the scientific community shed light onto the puzzling concept of consciousness.

With this in mind, Goldman and colleagues built the TVB-AdEx model, a multi-scale whole brain model that connects the behavior of individual neurons to whole brain dynamics [5, 13]. To build this model, one starts at the microscopic scale by building a spiking network containing excitatory and inhibitory Adaptive Exponential (AdEx) integrate and fire neuron models [14]. Then, the mesoscopic AdEx mean-field model is derived from this network, which captures the aggregated spiking dynamics of the network at the mesoscopic scale [15, 16]. The final step is to bridge from the mesoscopic to the macroscopic scale by forming a network of these AdEx mean-field models, connecting them using anatomical information. This multi-scale model is capable of exhibiting clear transitions from fast, low amplitude and complex (AI) to slow oscillating, high amplitude (Up and Down, UD) spontaneous activity when changing the adaptation value that simulates the change in acetylcholine concentration [10]. The model is also capable of reproducing different evoked patterns of activity for conscious and unconscious states[13], consistent with experiments [9, 17, 18].

The TVB-AdEx model is thus a high-dimensional dynamical model with a multitude of parameters that must be understood and estimated. Fortunately, many parameters can be informed through physiological and mathematical constraints [15, 16, 19]. However, the values of at least five parameters can still vary within a considerable range, allowing the model to explore many, possibly interesting, regimes. Nevertheless, the impact of these parameter values on the model has not been thoroughly studied yet because of the incredibly large amount of parameter combinations that can be generated.

Moreover, under certain conditions that were not yet thoroughly explored, the AdEx mean-field models that form the TVB-AdEx network can stabilize at a pathologically high level of activity. Simulations where this “paroxysmal” activity appears do not faithfully model neural dynamics associated with either healthy conscious or unconscious brain states, but rather seem to represent aberrant dynamics such as in epileptic states.

In this work, High Performance Computing (HPC) has been applied to perform a dense grid parameter exploration of the model. Thanks to this exploration, a characterization of the regions where the pathological activity is most probable has been carried out. Additionally, it has been found that the transitions between AI and UD when changing the adaptation level are remarkably robust. Finally, it has been possible to explore how functional patterns relate to anatomical constraints, a characteristic of the model resembling empirical results that had not been yet explored.

## 2. Methods

### 2.1. The TVB-AdEx Model

The TVB-AdEx model is a biologically informed, brainscale cortical model built using The Virtual Brain (TVB) platform. Using MRI scans, one can divide a scanned brain into different anatomical regions and estimate the strength of the connections between regions using tractography analyses [20]. From this data one can build the TVB-AdEx model as a network of AdEx mean-field models, each mean-field model describing the neuronal activity of one of the anatomical regions in the scanned brain. The strength of the interactions between the mean-field models are determined by the connectivity matrix, also called connectome, obtained from the MRI tractography analyses. The parcellation used for this model divides the brain into 68 regions (berlinSubjects/QL_20120814 from https://zenodo.org/record/4263723, [20]) and, therefore, 68 AdEx mean-field models make up the TVB-AdEx model.

Each one of the 68 AdEx mean-field models reproduces the mean behavior of a spiking network made up of 10^4^ Adaptive Exponential (AdEx) integrate and fire neurons (80% of them being excitatory and the other 20% inhibitory) [16]. Through a Master equation formalism [15], the mean-field model describes the general activity of the neural populations in the spiking network with seven differential equations [16, 19]. When spike-frequency adaptation is low (simulating conscious brain states with high levels of neuromodulation [5, 16]) the mean-field model is able to reproduce AI states through noisy perturbations around a stable non-zero fixed point (Up fixed point). When adaptation is increased (simulating reduced neuromodulation in the brain), the model also reproduces Up-Down states by cyclically traveling between the fixed point at the origin (Down fixed point) and the Up fixed point [16].

Note that there are two different networks needed to build this macroscopic model: the spiking network, modeling 10^4^ neurons at the microscale, and the TVB-AdEx network, consisting of the 68 inter-connected AdEx mean-field models. From now on, we will only refer to the spiking network when the term “spiking” is explicitly mentioned, in any other case, the word “network” will refer to the TVB-AdEx model.

The variables that describe the mean activity of the excitatory and inhibitory populations are the mean excitatory and inhibitory firing rate, *v*_*e*_ and *v*_*i*_, respectively. Under normal working conditions, both mean firing rates remain under 100 Hz, typically in the range [0, 50] Hz. However, the dynamical landscape of the AdEx mean-field model also contains a fixed point at a pathologically high level of activity, around 190 Hz, where neurons in the populations fire immediately after their refractory period (see Figure 10).

This fixed point does not disappear when building the TVB-AdEx network. Instead it becomes more difficult to detect under which conditions one, or multiple, mean-fields in the network stabilize at that pathological point, an event that we will refer to as “paroxysmal firing rates” or “paroxysmal fixed points”. We argue later that these paroxysms are more relevant to epileptic states, so they were avoided in this study.

In the TVB-AdEx model, the long-range connections between mean-field models are excitatory, as is shown in the left panel of Figure 1, and their strength is given by the structural connectivity matrix *C*_*jk*_ derived from tractography data, shown in the right panel of Figure 1.

**Figure 1:**
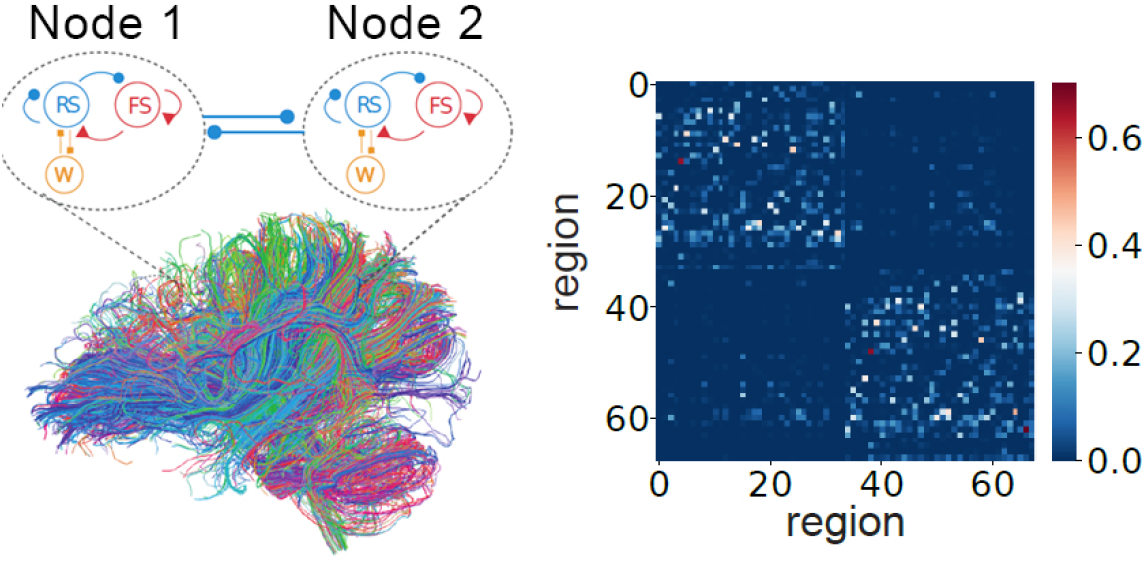
Left panel shows a diagram of two of 68 mean-field nodes in the simulation, along with tractography data informing long-range connections between mean-field nodes. Right panel shows the strength connectivity matrix, *C*_*j,k*_, between the 68 nodes. Reproduced with permission from [13].

Equation 1 describes node *k*’s evolution in time of the firing rate of the excitatory and inhibitory (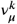 with *μ* = {*e, i*}) populations. Equation 2 describes the covariance between populations *λ* and *η* (*λ, η* = {*e, i*}) in node *k* and Equation 3 describes the evolution in time of the adaptation variable of node *k, W*^*k*^. Note that Einstein’s index summation convention is used, omitting summation symbols and summing over repeated indices.

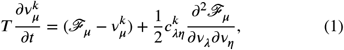

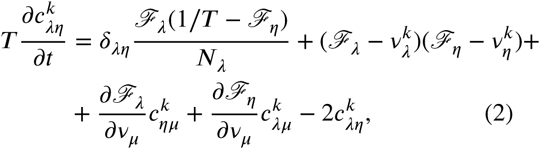

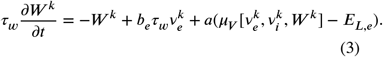

The parameter *T* sets the time scale of the mean-field models’ dynamics. Since the choice of *T* is delicate when dealing with non-stationary dynamics [16], it has been one of the parameters included in this work’s parameter sweep. The function 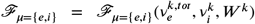 is the transfer function of the excitatory and inhibitory AdEx neurons in the spiking network, which determines the firing rate of the neuron when receiving 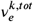 and 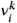 as inputs. These functions are obtained through a semi-an^*i*^alytical derivation which is explained in detail in [16].

Parameters *b*_*e*_ and *a* come from the AdEx single neuron model, at the microscopic scale, and stand for spike-triggered adaptation (in pA) and subthreshold adaptation (in nS), respectively. The value of *a* has been fixed to *a* = 0 nS, while different values of parameter *b*_*e*_ will be explored during this work as we are interested in observing how the TVB-AdEx model’s activity changes when spike-triggered adaptation varies. 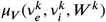 is a function that returns the average membrane potential of the population, obtained when deriving the mean-field model [16]. The *E*_*L,e*_ parameter is also related to the single neuron model used in the spiking network. It stands for the leakage reversal potential of AdEx excitatory neurons (their resting membrane potential [14]) and its impact on the model is discussed in this work. Although it is not explicitly present in the equations, the leakage reversal potential of inhibitory neurons *E*_*L,i*_ also has an important effect on the outcome of the model through both *μ*_*V*_ and the transfer functions, so it has also been included in the parameter sweep.

Finally, 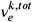 corresponds to the total incoming excitatory input to a neuronal population given by:

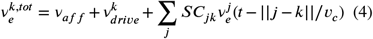

where *v*_*aff*_ is an afferent transient input, 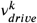 is an external noise simulated by an Orsntein-Uhlenbeck process, *C*_*jk*_ is the connectivity between regions *j* and *k* (with *C*_*kk*_ = 1) and 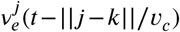 is the activity of the excitatory population in node *j* with a delay corresponding to the distance between the regions divided by the axonal propagation speed *v*_*c*_. The *S* parameter is phenomenologically tuned to modify the overall inter-region connectivity strength and, therefore, it is the final parameter included in this work’s analyses.

### 2.2. Parameter Exploration

As previously mentioned, such a complex model contains many parameters that need to be understood and to have reasonable, physiological ranges determined for them. Most of those variables can be set based on biological or mathematical arguments, but there remains a subset of parameters whose impact needs to be studied to have a deeper and general understanding of the model. In Table 1, one can find the characteristics of the parameters chosen and the reason for their choice.

**Table 1.** Name, description, reason of choice, range and units of the parameters chosen for the parameter sweep.

| Parameter | Description | Reason of choice | Range | Units |
| --- | --- | --- | --- | --- |
| $S$ | Coupling strength between nodes. | Has to be chosen phenomenologically. | [0, 0.5] | Dimensionless |
| $E_{L,i}$ | Leakage reversal potential of AdEx inhibitory neurons. | Resting membrane potential of a neuron might vary depending on external conditions. | [-80, -60] | mV |
| $E_{L,e}$ | Leakage reversal potential of AdEx excitatory neurons. | Resting membrane potential of a neuron might vary depending on external conditions. | [-80, -60] | mV |
| $T$ | Timescale of the AdEx mean field model. | Has to be chosen phenomenologically. | [5, 40] | ms |
| $b_e$ | Adaptation strength of excitatory AdEx neurons. | Models the change in neuromodulation that induces transition between AI and UD. | [0, 120] | pA |

For each parameter, 16 evenly spaced values are obtained inside the described range. A simulation would have been run for each of the possible combinations of parameter values, which would result in having to analyse 16^5^ differently parametrized TVB-AdEx configurations. However, preliminary results showed that neuronal activity remains silent when *E*_*L,i*_ is significantly greater than *E*_*L,e*_, a result of the inhibitory populations being more active than the excitatory ones. For this reason, only those combinations where *E*_*L,i*_ < *E*_*L,e*_ + 4 mV have been simulated. In the end, a total number of 675,840 different configurations have been analyzed.

### 2.3. High Performance Computing

Though more computationally tractable than simulating a brain-scale network of spiking neurons, simulating the TVB-AdEx consisting of mean-field populations remains a relatively computationally expensive process, taking approximately a minute to simulate one second of activity on a personal computer. For each parameter combination that is going to be studied, at least five seconds of activity need to be simulated. The analyses of the simulation might take up to an extra minute to be executed. Fortunately, the simulations needed to perform a parameter sweep can be easily parallelized.

For these reasons, the use of HPC has made this parameter sweep possible. We ran our scripts on the JUSUF supercomputer in the Jülich Supercomputing Centre [21], which consists of 187 nodes, with each node having two AMD EPYC 7742 @2.25 GHz processors for a total of 128 cores, 256GB DDR4 of RAM and 1TB NVMe for memory [22].

A profiling of a five-seconds TVB-AdEx simulation, together with the corresponding feature extraction pipeline, showed that running the script required less than 600 MB of RAM at any point in time, less than one GB of static memory for I/O operations, and approximately 6 minutes of execution time (see Figure 8 in the Supplementary Information). Thus, it was possible to run 128 simultaneous simulations on each JUSUF node, increasing significantly the simulation and analysis speed.

### 2.4. Feature Extraction on Spontaneous Activity

In order to visualize the results of the parameter exploration, a feature extraction pipeline has been applied, obtaining a series of metrics (or features) that represent the overall behavior of the TVB-AdEx model for each parameter combination studied (see Figure 9 in the Supplementary Information for a schematic representation).

For each parameter combination, a five seconds simulation of the TVB-AdEx model has been run with a time step of 0.1 ms. The first two seconds following initialization of the simulation are considered a transient state and, therefore, ignored. Afterwards, the array of data containing the evolution in time of the 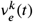 of the 68 regions of the model is recovered to apply the feature extraction.

In order to analyze the spectral characteristics of a parameter combination, the Fourier Transform is applied to each of the 68 time series in the 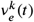 array, obtaining a 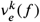 array in the Fourier space. Then, each element of the array is squared and an average over the nodes, obtaining a single *PSD*(*f*) curve representing the average Power Spectral Density (PSD) of the nodes in the network, from which spectral features will be obtained.

Table 2 displays the features that will be analyzed in this report, along with a short description of how they are obtained. Other features that have been computed but whose analyses are too extensive for the limited space of this report are displayed in Table 3 in the Supplementary Information. As a final note, the excitatory firing rate has been is the source of the features shown in this manuscript, since there is little observable difference between features obtained with *v*_*e*_ and *v*_*i*_.

**Table 2.**
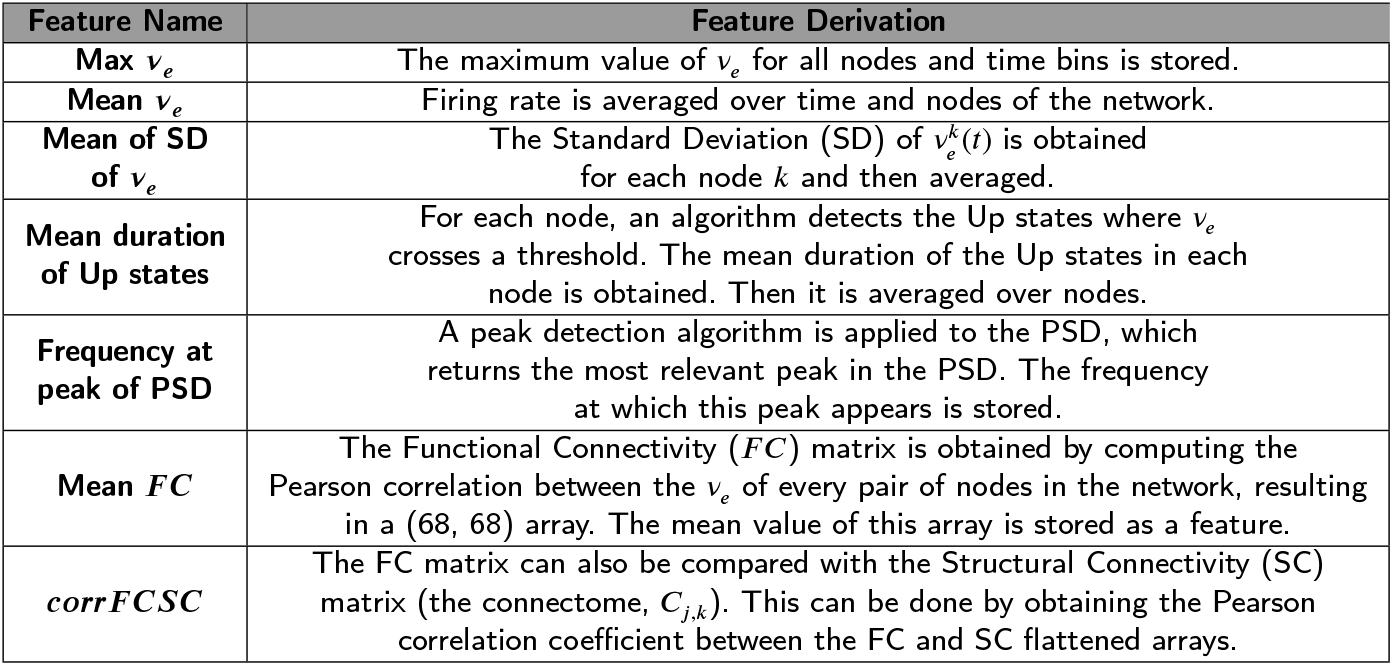
Table containing the features that will be analyzed in this work, along with a short description of their derivation.

**Table 3.** Remaining features that have been obtained for each parameter combination but that have not been included in the main text’s analyses. It is expected to make use of them in further work.

| Feature Name | Feature Derivation |
| --- | --- |
| Mean duration of Down states | For each node, an algorithm detects the Up states where $v_e$ crosses a threshold. A Down state is the time between the moment when $v_e$ becomes smaller than the threshold and crosses it again. The mean duration of Down states in each node is obtained. Then it is averaged over nodes. |
| Mean PLI value | The average Phase Locking Index (PLI) of the whole network is obtained by averaging the PLI matrix, result of computing the PLI value for each pair of nodes in the network. See Appendix of [10]. |
| Power Law of PSD | The PSD of neuronal activity is expected to follow a Power Law of the type $PSD(f) = \frac{b}{f^a}$. In a loglog plot, this is equivalent to fitting a linear regression, from which we store the $a$ value. |
| Score of PSD Power Law | The r-score obtained when fitting the linear regression on the loglog plot of the PSD is stored. |
| Relative Powers of frequency bands | 5 bands are defined in the (0.5, 100) Hz range of frequency:<br>$\delta \in (0.5, 4)$ Hz, $\theta \in (4, 8)$ Hz,<br>$\alpha \in (8, 12)$ Hz, $\beta \in (12, 30)$ Hz, and $\gamma \in (30, 100)$ Hz.<br>The $\int_{B_i}^{B_f} PSD(f)df$ gives the spectral power in a band.<br>Dividing by the total power (integrated between 0.5 and 100 Hz) we obtain the fraction of spectral power in a band. |
| *corrFCTL* | Similar to the *corrFCSC*, the Pearson correlation between the flattened Functional Connectivity Tract Lengths (TL) matrices is obtained. The TL matrix indicates the length of the anatomical tracts of white matter between two regions in the whole brain parcellation. |
| Ratio top z-score of DMN | The Default Mode Network (DMN) are a set of regions in the brain that are correlated during resting brain dynamics (when the brain is not performing any cognitive task). It has also been shown that its elements remain correlated when falling asleep. It is also believed that it is strongly tied to anatomical pathways [25, 26, 27].<br>For these reasons, since one of the strengths of the model is its anatomically informed connectivity, correlations between regions associated to the DMN were studied. The regions considered as the DMN for the given parcellation can be seen in the Figure immediately below. As done in fMRI studies, one of the regions (left precuneus) is taken as a seed and the correlations between the seed and all the other regions are obtained (Fisher transformation is applied to obtain a zscore). Then, the resulting metric is obtained by calculating how many of the DMN components appear in the top ten most correlated anatomical regions with the seed. One would expect that most of the DMN components appear in this top ten, however, results have not been conclusive. |

## 3. Results

### 3.1. Normal and paroxysmal states

First of all, it is fundamental to have a precise characterization of when the dynamics of the TVB-AdEx model exhibit firing rates in the physiological range or if they exhibit paroxysmal or aberrant activity. In this case, one, or multiple, mean-field model reaches the pathologically high activity fixed point. Since this pathological fixed point is found at around 190 Hz, well above the “normal” working range, it is possible to analyze the occurrence of this event by studying the maximum *v*_*e*_ reached for every one of the 675,840 different configurations: if the maximum value of *v*_*e*_ exceeds 175 Hz (activity only reaches such high values in the paroxysmal fixed point), that parameter configuration is counted as exhibiting paroxysmal dynamics.

The parameters that mostly favor the emergence of paroxysmal dynamics are *S, E*_*L,i*_ and *E*_*L,e*_ (see Figure 11 in the Supplementary Information), which are all related to the level of excitatory/inhibitory balance in the network, consistent with previous results [23].

It is thus possible to build a 3D histogram, with the objective of seeing how these three different parameters interact between them to favor pathological activity, shown in Figure 2. This is done by fixing the values of *S, E*_*L,i*_ and *E*_*L,e*_ and counting how many Paroxysmal Fixed Points (PFPs) appear when varying the remaining parameters (*T* and *b*_*e*_).

**Figure 2:**
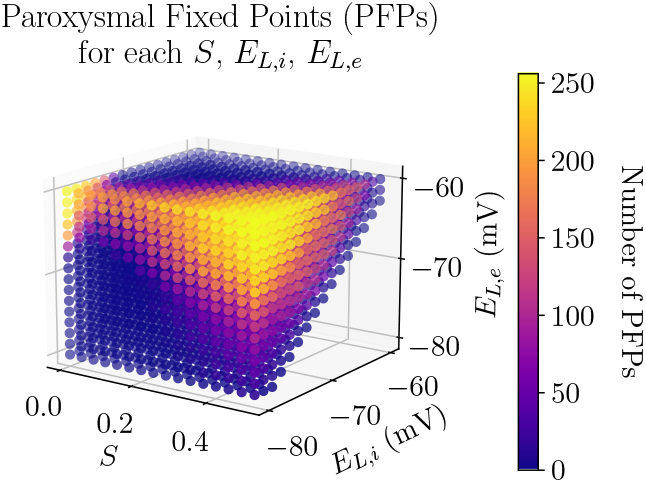
Number of Paroxysmal Fixed Points (PFPs) as a function of *S, E*_*L,i*_ and *E*_*L,e*_ parameter values.

As one could expect, the probability of reaching a PFP is largest in the region of highest *S* and *E*_*L,e*_ and lowest *E*_*L,i*_, since it groups together all the conditions that favor the appearance of PFPs. Curiously, one can also see that the *S* = 0, high *E*_*L,e*_, low *E*_*L,i*_ corner also exhibits higher probability of exhibiting paroxysmal activity, indicating that disconnected mean-field modes tend to reach this pathological activity more easily. It seems that near the *E*_*L,i*_ = *E*_*L,e*_ diagonal is a relatively safe working region, which we will use in the following section.

### 3.2. Robustness of AI to UD Transition When Increasing Adaptation Strength

The parameter sweep described in this work was designed to initially study the robustness of transitions between AI and UD states when changing the spike-frequency adaptation strength, modeled by the *b*_*e*_ parameter. The main objective was to determine the regions of the parameter space where this transition takes place, as well as to look for other possible dynamics of the system that might relate to empirical findings. Therefore, a representative feature of the transition between AI and UD states (the *v*_*e*_ standard deviation) is shown in Figures 3 and 4 for two different regions of the parameter space, together with the time evolution of the TVB-AdEx network’s excitatory and inhibitory firing rates. In the TVB-AdEx, AI states are described by small amplitude stochastic perturbations around a fixed point of sustained activity while UD states are described by an oscillatory traveling between the origin (Down state) and this AI fixed point (Up state). Thus, AI states are characterized by higher mean *v*_*e*_, since there are no silent periods; smaller standard deviations, due to their range being narrower; and much longer Up state duration than UD states, since they are at a constant Up state.

**Figure 3:**
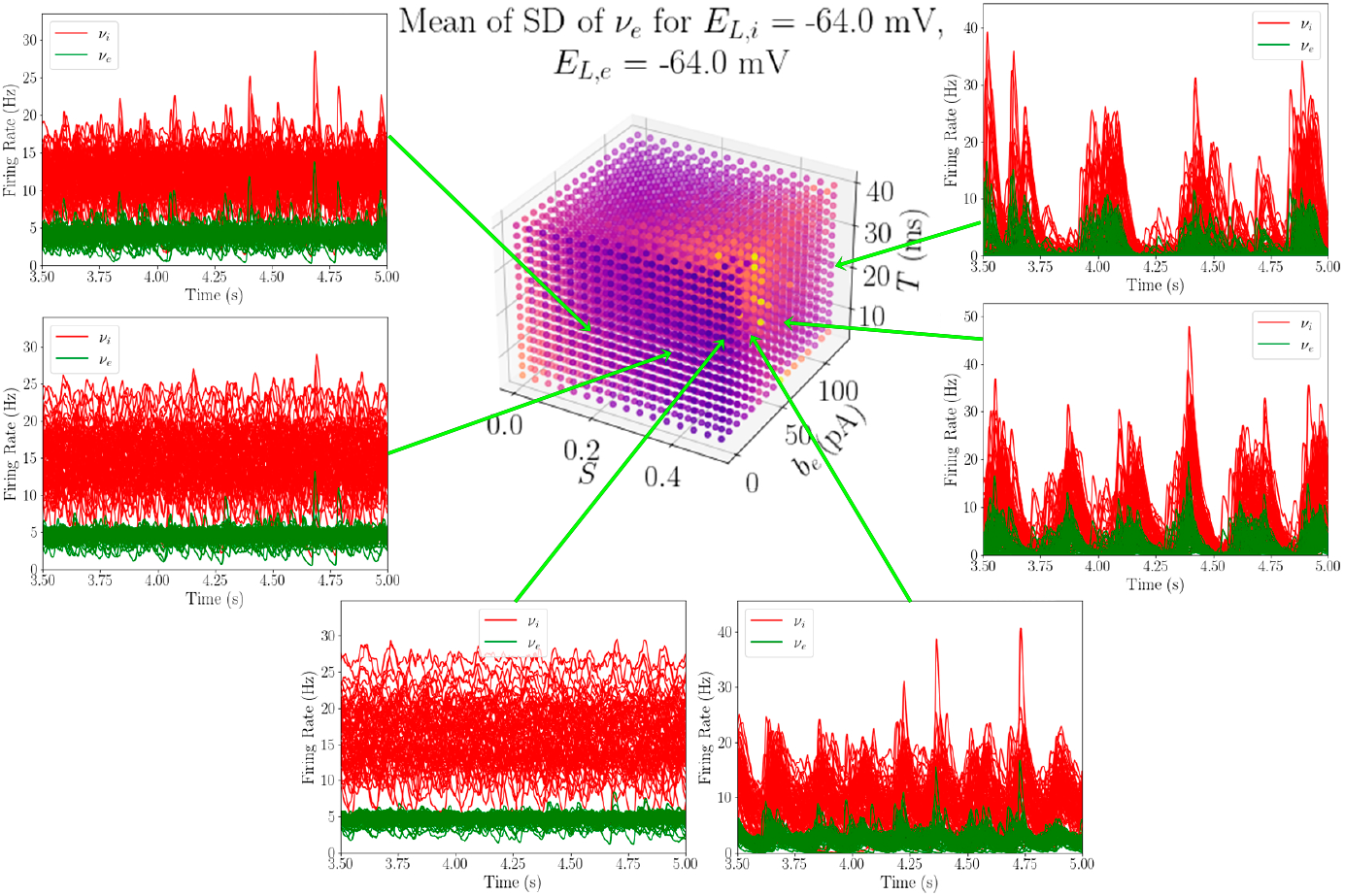
Mean standard deviation values in a subspace of the *depolarized region* of the parameter space. In the floating panels, the time evolution curves of the inhibitory and excitatory populations of the 68 AdEx mean-field models are plotted for six different parameter combinations in the depolarized region (*E*_*L,i*_ = *E*_*L,e*_ = −64 mV) during their last one and a half seconds. In *b*_*e*_ = 0 pA, for intermediate values of *S*, one can see how AI states appear, consistent with the low value of SD shown in the feature plot. For *S* = 0.5, one can see the transition between AI and UD states when increasing *b*_*e*_.

**Figure 4:**
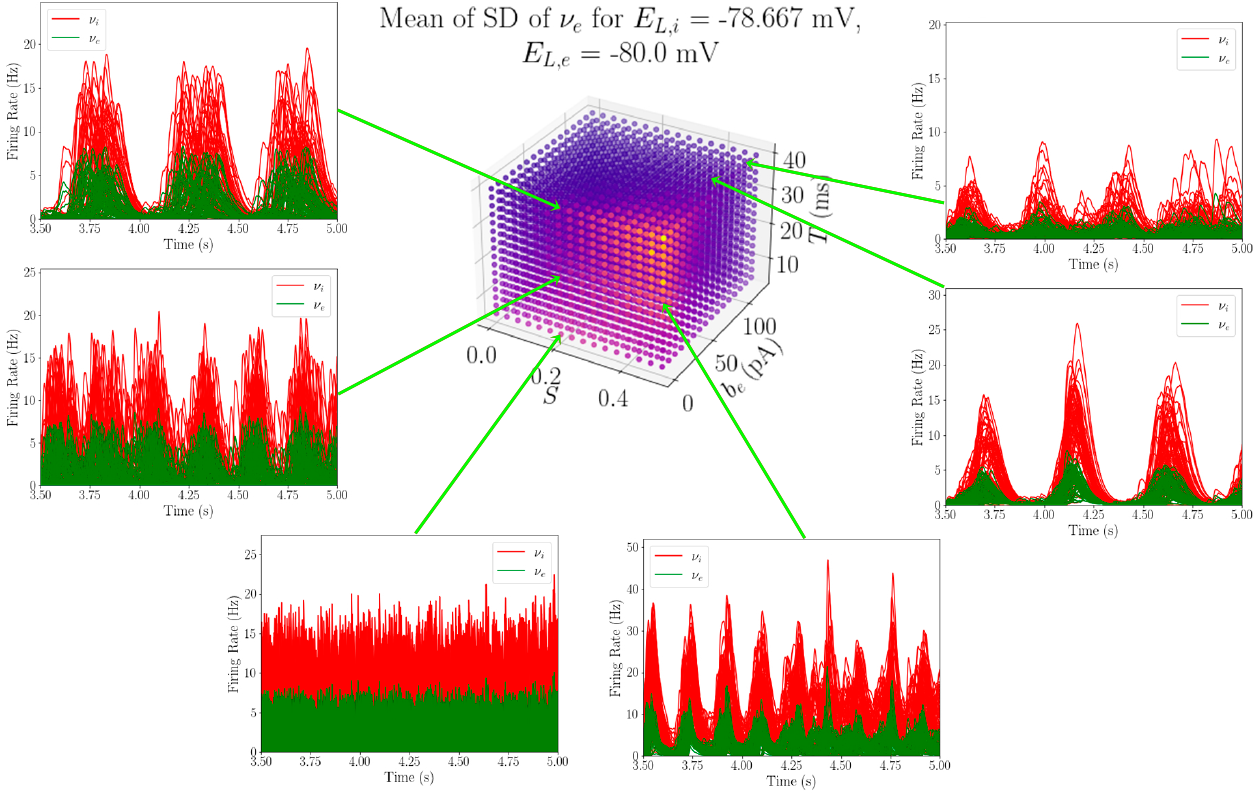
Mean standard deviation values in a subspace of the *hyperpolarized region* of the parameter space. In the floating panels, the results of running the TVB-AdEx model for six other parameter combinations are plotted. In the hyperpolarized region (*E*_*L,i*_ = −78.667 mV, *E*_*L,e*_ = −80 mV) one can clearly see the effect that the leakage reversal potential values have on the behavior of the model. Here, AI states do not appear even for *b*_*e*_ = 0 pA and high values of *S*. Instead, this sub-region of the parameter space is dominated by UD dynamics. For small *T* values, one can see how the dynamics are very fast and desynchronized, making it hard to visualize. However, seeing that the excitatory firing rates reach values of silence, one can conclude that they consist of faster UD states.

Figure 3 shows the standard deviation (averaged over time and nodes of the network) of *v*_*e*_ when fixing *E*_*L,i*_ and *E*_*L,e*_ at equal and relatively high values (−64 mV). We will refer to the region of the parameter space where −60 mV ≥ *E*_*L,e*_ ≥ −65 mV as the *depolarized region*. Additionally, the time evolution of the excitatory (green) and inhibitory (red) firing rates are shown for different parameter combinations of the model (each panel corresponding to a simulation computed with the parameters of a point in the cube).

One can clearly see that, for low *b*_*e*_ and for most *S* and *T* values, AI-like dynamics are found (three left-most panels). These patterns of activity are characterized by high mean *v*_*e*_, low standard deviation, high duration of Up states and higher frequencies of *PSD* peak, which are plotted in Figure 12 in the Supplementary Information. When increasing the spike-frequency adaptation, one can see a sharp change in the behavior (three right-most panels), towards an UD-like behavior where the standard deviation of the firing rates increases, while the mean firing rate decreases, together with the Up-state duration and the frequency at the peak of the *PSD* (again, shown in Figure 12).

It is also interesting to explore the influence of these same three parameters but in a much different region. In Figure 4, similar plots to those in Figure 3 are shown, but in this case, the parameters *E*_*L,i*_ and *E*_*L,e*_ are fixed to *E*_*L,i*_ = −78.667 mV and *E*_*L,e*_ = −80 mV, while *S, b*_*e*_ and *T* are swept in the same range. In this situation, the inhibitory and excitatory populations are at the lower limit of the physiologically relevant leakage reversal potentials values (inhibitory populations are slightly less hyperpolarized for plotting purposes, to avoid some of the parosysms that may appear when *E*_*L,i*_ = −80 mV). Thus, we will refer to the region where −75 mV ≥ *E*_*L,e*_ ≥ −80 mV as the *hyperpolarized region*.

Now, the standard deviation values show a non-expected peculiar behavior: the nodes of the TVB-AdEx network exhibit the highest standard deviation of *v*_*e*_ for the lowest values of *b*_*e*_, which should, in theory, correspond to AI states, associated to low standard deviation. Looking at the time evolution curves of the selected parameter combinations, it is clear that these standard deviation values come from the appearance of UD states, even when there is no adaptation in the system. This type of behavior could be associated to anesthetized brains, a state of unconsciousness where neurons are typically hyperpolarized [24].

This modification of the behavior when *E*_*L,i*_ and *E*_*L,e*_ shift between the *depolarized* and the *hyperpolarized* states is relatively smooth when increasing the leakage reversal potential of the populations (increasing *E*_*L,i*_ = *E*_*L,e*_ = *E*_*L*_ from −80 to −60 mV), as can be seen in Figure 5. When there is no spike frequency adaptation, if the neuronal populations are hyperpolarized, UD states dominate the dynamics. However, when the neurons reach their typical *E*_*L,e*_ and *E*_*L,i*_ values, AI states start to appear.

**Figure 5:**
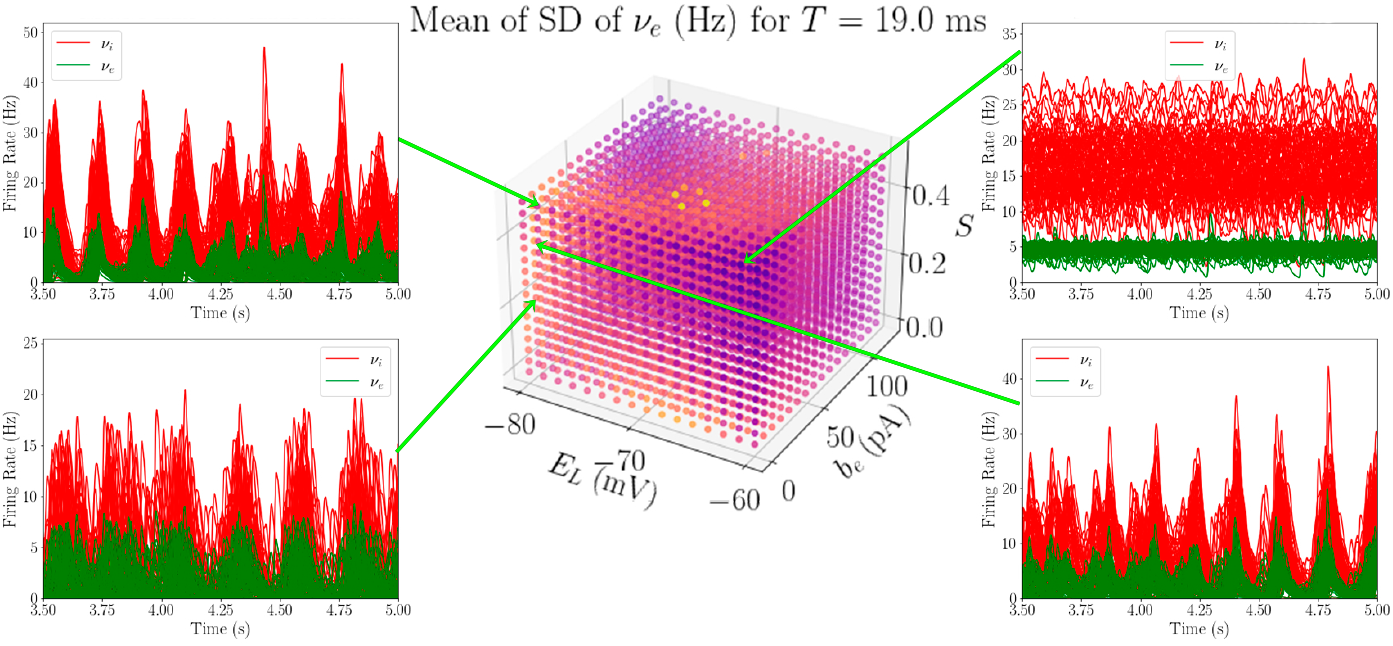
Mean standard deviation values when fixing *T* = 19*ms* and exploring *E*_*L,i*_ = *E*_*L,e*_ = *E*_*L*_, *b*_*e*_ and *S*. When fixing adaptation to *b* = 0*pA* but changing *E*_*L*_, there is a clear transition between AI (higher values of *E*_*L*_) and UD states (low values of *E*_*L*_), which could be associated to a descent from consciousness towards anesthetized states [24].

Thus, this parameter sweep has shown a remarkable robustness of the transition between AI and UD states when varying the adaptation strength, *b*_*e*_, especially in the *depolarized region* of the parameter space. Additionally, a transition from AI behavior to slow-wave oscillations has been found when hyperpolarizing the neural populations in the model, which could be related to transitions between conscious and anesthetized states.

### 3.3. Relationship Between FC and SC

The parameters used in the original TVB-AdEx model in [13] show a transition from asynchronous AI to synchronized UD states, consistent with LFP empirical recordings [17]. In order to extend that line of work, one of the objectives of this project was to investigate whether transitions from low to high synchronization between network nodes with increasing adaptation is maintained throughout the studied parameter space. Additionally, empirical studies show that the Structural Connectivity (SC), described here by the matrix *C*_*j,k*_, tends to be the substrate upon which *FC* correlations appear during unconscious states, while the repertoire of *FC* patterns varies considerably and diverges from the SC in conscious states [7, 8, 25]. So, the relationship between the FC patterns and the SC of the TVB-AdEx model has been studied when modeling a descent towards modeled unconscious dynamics by increasing *b*_*e*_.

To do that, the evolution of the mean value of the FC matrix, (from now on *FC*) and the Pearson correlation between the FC and SC matrix (from now on *corrFCSC*), described in Table 2, are studied as a function of *b*_*e*_ for each combination of *S, E*_*L,i*_, *E*_*L,e*_ and *T* values. In other words, parameters *S, E*_*L,i*_, *E*_*L,e*_ and *T* are fixed and one observes how *FC* and *corrFCSC* vary when increasing *b*_*e*_ from 0 to 120 pA, obtaining the *FC*(*b*_*e*_) and *corrFCSC*(*b*_*e*_) traces for that parameter combination. This results in 42,240 different *FC*(*b*_*e*_) and *corrFCSC*(*b*_*e*_) traces that need to be analyzed. To approach this cumbersome task, the *FC*(*b*_*e*_) and *corrFCSC*(*b*_*e*_) traces have been clustered (using a K-means clustering algorithm) with the objective of automatically grouping them in six *FC*(*b*_*e*_) classes and six *corrFCSC*(*b*_*e*_) classes. Those *S, E*_*L,i*_, *E*_*L,e*_ and *T* combinations containing paroxysmal activity at any point of *b*_*e*_ were ignored from this analysis. In Figure 6, one can see the centroids of the six *FC*(*b*_*e*_) and *corrFCSC*(*b*_*e*_) classes. In K-means clustering, the centroid of a class represents the average behavior of all the traces associated to that class, giving a general idea of their tendency.

**Figure 6:**
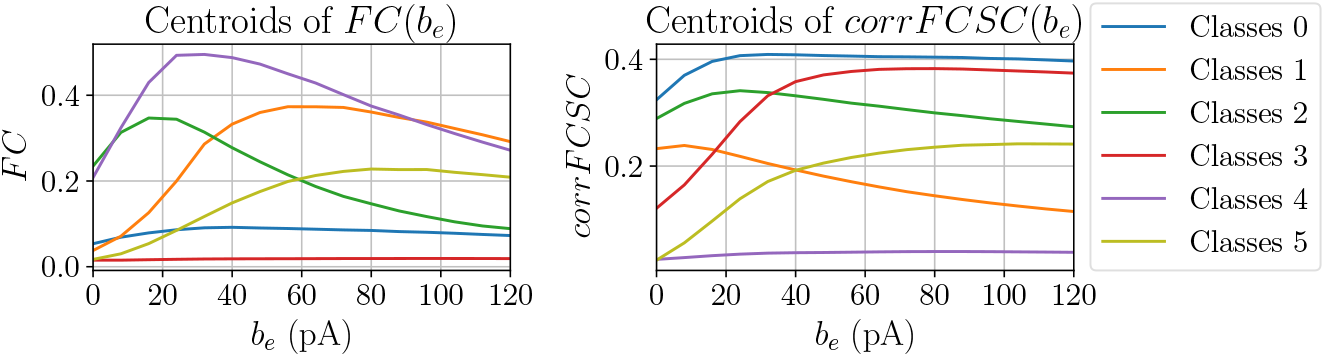
Centroids of the *FC*(*b*_*e*_) and *corrFCSC*(*b*_*e*_) classes, determined by a K-means algorithm with six clusters. Note that colors are used to distinguish between classes in the same category, *FC*(*b*_*e*_) and *corrFCSC*(*b*_*e*_) classes are not related, a priori, although they share the same colors. The number of classes has been chosen for plotting purposes.

The left panels in Figure 7 shows to what *corrFCSC*(*b*_*e*_) class each parameter combination is associated to in some parts of the *depolarized region* of the parameter space, while the right panels display the fraction of times that one of the parameter combinations in the left panel is assigned both *FC*(*b*_*e*_) class *i* and *corrFCSC*(*b*_*e*_) class *j*.

**Figure 7:**
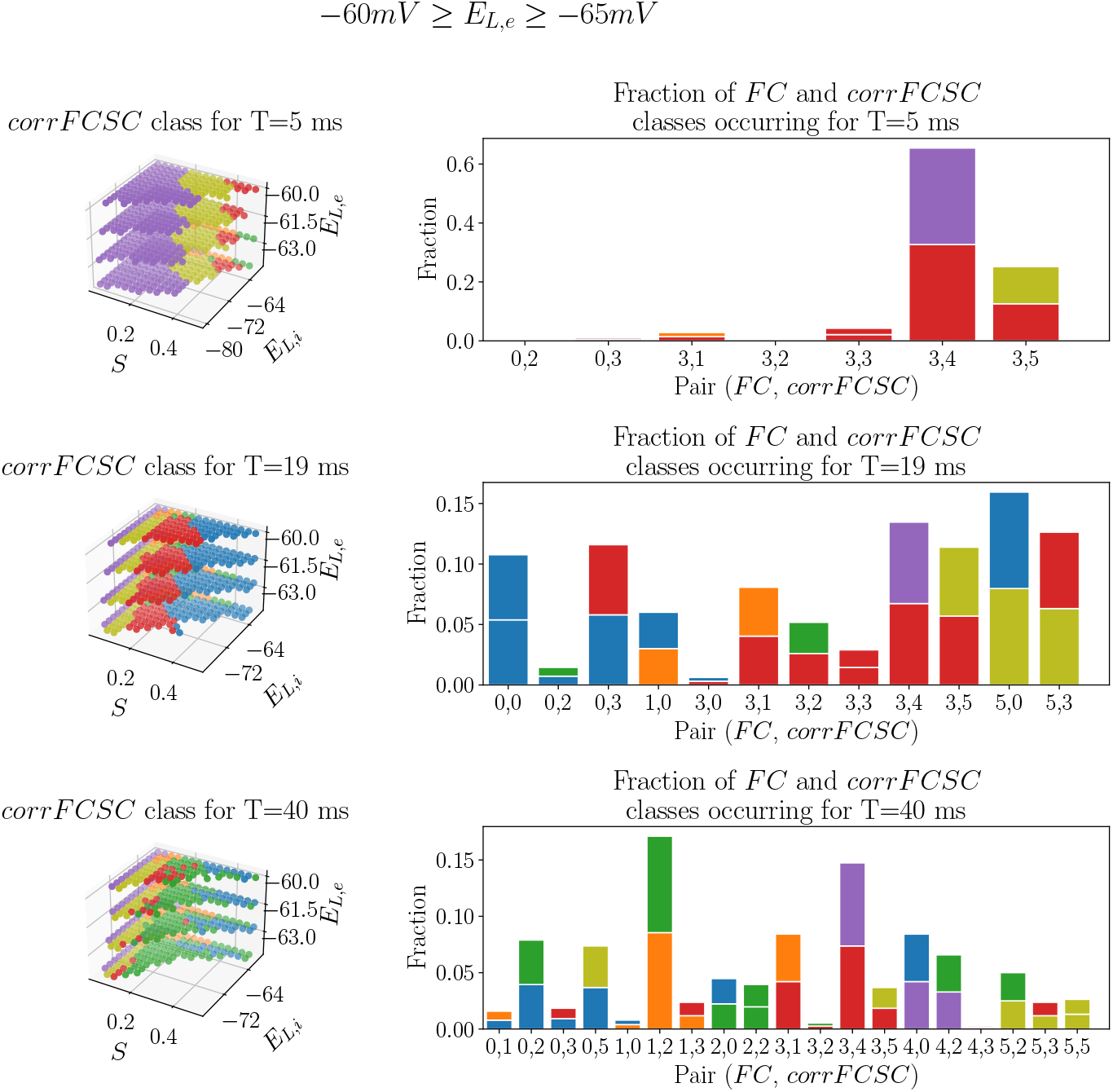
Distribution of *corrFCSC*(*b*_*e*_) classes as a function of *S, E*_*L,i*_ and *E*_*L,e*_ values for different *T* values (left panels) and histograms counting the fraction of times a point in the parameter space is assigned to *FC*(*b*_*e*_) class *i* and *corrFCSC*(*b*_*e*_) class *j* (right panels) for the *depolarized* range of *E*_*L,e*_ (−60 mV ≥ *E*_*L,e*_ ≥ −65 mV). The bottom color of the histograms’ bars are the same as the pair’s *FC*(*b*_*e*_) color, while the top color is the same as the pair’s *corrFCSC*(*b*_*e*_) color.

One can see that a low value of *T* = 5*ms* results in a predominance of little to no increase of *corrFCSC* with *b*_*e*_, being purple the most frequent class (recall *corrFCSC*(*b*_*e*_) centroids from Figure 6). Also, looking at the histogram, one can see that most *corrFCSC*(*b*_*e*_) classes are paired with the red *FC*(*b*_*e*_) class, implying that *FC* remains extremely low throughout the whole *b*_*e*_ range for *T* = 5 ms. For *T* = 19 ms, both blue, red and olive *corrFCSC*(*b*_*e*_) classes appear quite frequently. Analysing the centroids in Figure 6, blue, red and olive *corrFCSC*(*b*_*e*_) classes are the ones that show an increase in correlation when increasing *b*_*e*_, consistent with empirical recordings [7, 8, 25]. Examining the histograms, one can see that the blue, red and olive *corrFCSC*(*b*_*e*_) are also paired to an interesting variety of *FC*(*b*_*e*_) classes. Finally, for *T* = 40 ms, most of the points in this parameter region are assigned to the green *corrFCSC*(*b*_*e*_) class, which is characterized by a peak and slight decrease of *corrFCSC* when increasing adaptation. Although this region is the most heterogeneous of the three in terms of different class pairs, the most significant *corrFCSC*(*b*_*e*_) classes (blue, red and olive) are scarce.

The behavior of the *hyperpolarized region* is considerably different (see Figure 14 in the Supplementary Information). In the *hyperpolarized region*, for *T* = 19 ms, blue and green *corrFCSC*(*b*_*e*_) points, whose centroids start at a considerably high level of *corrFCSC*, dominate the region. This is consistent with the fact that, in the *hyperpolarized region*, one can find UD states associated to unconscious dynamics even for low values of *b*_*e*_.

Thus, these results show that during slow-wave activity, the FC patterns of the TVB-AdEx model typically become more constrained by the anatomical connectivity than during conscious-like states, which is in close agreement with the empirical results obtained in [7, 8, 25], especially for intermediate values of *T*, such as *T* = 19 ms.

## 4. Discussion and Future Perspectives

In this work, HPC resources have been used to analyze the spontaneous and evoked activity of the TVBAdEx whole brain model throughout a large parameter space. It has been found that transitions between conscious and unconscious-like dynamics when increasing adaptation strength *b*_*e*_ are largely robust, especially when the leakage reversal potential of excitatory populations is −60 mV ≥ *E*_*L,e*_ ≥ −65 mV. Moreover, it has been shown that it is possible to obtain slow-wave activity, associated to unconsciousness, when strongly hyperpolarizing the neural populations of the model via *E*_*L,i*_ and *E*_*L,e*_, independently of the level of spike-frequency adaptation. Finally, it has been found that the FC patterns arising from unconscious-like dynamics are usually closer to the underlying anatomical connectivity than those from conscious-like states, which is consistent with results from several empirical studies [7, 8, 25].

From studying the Paroxysmal Fixed Point probability in the TVB-AdEx model, one finds that the total number of PFPs in the parameter sweep is considerable, around 18%. The parameters that most strongly modulate the probability of reaching a PFP are *S, E*_*L,e*_ and *E*_*L,i*_, which are able to modify the balance between excitation and inhibition in the neural populations. This high number of PFPs could be decreased by applying a stronger long-range excitatory coupling on inhibitory populations with respect to excitatory populations [26], to compensate for the imbalance between excitation over inhibition that causes the pathological activity. This asymmetry could be further investigated to relate values of *E*_*L,e*_ and *E*_*L,i*_ to experimental findings of resting membrane potentials in different areas of the cortex. Because these paroxysmal points exhibit excessively high firing rates, and appear generally when there is an excess of excitation, they presumably represent epileptic-type activity, although this point would require a specific model to integrate biologically-realistic features of epilepsy, such as extracellular K^+^ dynamics [27].

The timescale of the model, expressed through parameter *T* has a powerful influence on the dynamics of the TVB-AdEx model. Apart from strongly modulating the frequency of oscillations of neuronal activity (clearly seen in Figures 12 and 13), it affects the overall level of synchronization and how the FC patterns relate to the anatomical connectivity matrix (Figures 7 and 14).

In the TVB-AdEx network, choosing low values of *T*, such as *T* = 5 ms, might result in violating one of the necessary assumptions used to build the AdEx mean-field model: that the system is memoryless over a certain time scale *T* [15]. This violation results in aberrant, noisy and fast oscillating activity (an example of this has been shown in Figure 4’s bottom-left panel). Therefore, correlations between nodes become extremely rare, resulting in low levels of mean FC and *corrFCSC* independently of the adaptation strength value.

For large values of the timescale parameter, such as *T* = 40 ms, the dynamics of the model slow down considerably, and correlations between regions increase, especially in UD states. This increase in correlations results in an almost homogeneous FC matrix, quite different from the SC matrix, which explains the overall decrease in *corrFCSC* as a function of *b*_*e*_ for *T* = 40 ms. Additionally, for large values of *T*, the memoryless assumption holds but inputs arriving to the mean-field models of the network at frequencies faster than 1/*T* (around 25 Hz for *T* = 40 ms) appear as a constant external drive [16].

Therefore, the results obtained for intermediate values of *T*, such as *T* = 19ms, support the idea that it is necessary to use a timescale slow enough that it does not violate the memoryless assumption but as fast as possible so that the loss of high-frequency information is kept to a minimum.

Even if it has been shown that the TVB-AdEx model can mimic both conscious and unconscious-like dynamics for many different parameter combinations, the model still has several noteworthy limitations.

First of all, it is a purely cortical model, which does not take into account the dynamics of subcortical structures such as the thalamus, which might be an interesting element to consider when modeling conscious, unconscious or anesthetized brain states [24, 28]. The TVB-AdEx model is able to reproduce both conscious and unconscious-like dynamics by describing only cortical activity and simplifying subcortical effects on the cortex via the transient and noisy inputs *v*_*aff*_ and 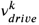, respectively, from Equation 4. Therefore, a brain-scale thalamo-cortical TVB-AdEX model might be extremely useful to understand the role of interactions between cortical and subcortical structures during conscious and unconscious states.

Additionally, this brain-scale cortical model assumes that each region shares the same parameters as the others. For instance, one could try to make use of an acetylcholine receptor density map to obtain a heterogeneous distribution of *b*_*e*_ parameters, similarly to what is done in [29].

Finally, most empirical functional connectivity studies nowadays make use of fMRI data, which describes neuronal activity at long temporal scales, with a typical sampling time of two seconds, while the sampling time of the TVB-AdEx model is of 0.1 ms, describing much faster neuronal dynamics. Bridging the gap between these two scales is out of the scope of this manuscript and will be explored in later work, possibly through the use of the BOLD monitor from the TVB library. Additionally, in the TVB-AdEx model, there is typically an increase of the mean FC value with increased adaptation, as has been shown in Section 3.3. However, this is not the case in fMRI signals, where the mean FC value decreases during the descent to deep sleep [25], a discrepancy that might be also explained by the difference in time scales and that should be further investigated.

For these reasons, reducing the computational cost of long simulations of the TVB-AdEx model would be extremely beneficial as it would allow to use the BOLD monitor from the TVB tool set to simulate BOLD signals. Analyzing simulated BOLD signals would allow to better relate the model’s behavior to empirical fMRI findings. Also, it would be possible to repeat the parameter sweep but studying the network properties of the FC matrix, in order to determine whether the TVB-AdEx model can also simulate known FC characteristics of brain pathologies [30, 31, 32] that have been analyzed from fMRI recordings. One possibility to speed up the simulations could be to make use of RateML, a tool that allows the user to generate accelerated CPU and GPU-ready TVB models [33].

Together, these results show that the TVB-AdEx model is a flexible and robust multi-scale model where changes on parameters that reflect microscopic processes induce multiple relevant macroscopic changes on both spontaneous and evoked activity, making it a viable model for studying different conscious and unconscious states from cells to whole-brain networks.

## Acknowledgments

Research supported by the CNRS and the European Union (Human Brain Project H2020-785907, H2020-945539). We thank Kevin Ancourt, Thierry Bal, Frederic Chavane, Nima Dehghani, Matteo di Volo, Sami El Boustani, Nicolas Hô, Trang-Anh E. Nghiem, Denis Paré and Yann Zerlaut for support and collaboration. We want to thank Sandra Diaz and Michiel Van der Vlag for their invaluable help in executing the simulations at the JUSUF supercomputer.

## Code Availability

The code used for the simulations presented in this paper can be found in https://github.com/davidaquilue/TVBAdEx_ParSweep and will be available online and in EBRAINS after publication.

## SUPPLEMENTARY INFORMATION

### A. Supplementary Methods

#### A.1. Profiling for HPC computing

**Figure 8:**
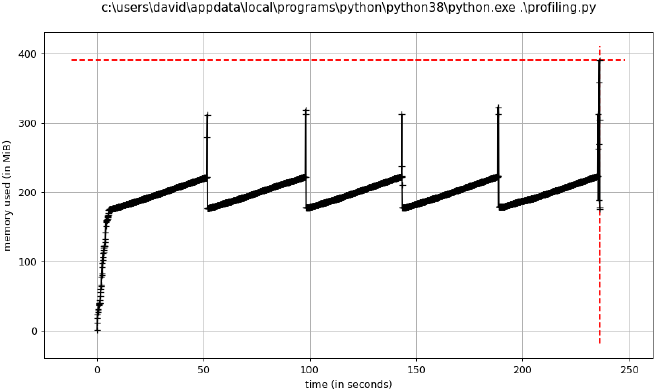
Memory and time consumption profiling for a single TVB-AdEx model simulation, without feature extraction. One can see that the amount of Random Access Memory (RAM) needed at each point never exceeds 400 MB. The feature extraction pipeline makes sure to only use the necessary resources, never surpassing the 600 MB limit. Thus, it is possible to exploit the 128 cores of a JUSUF node without worrying about exceeding the 256 GB RAM limit.

**Figure 9:**
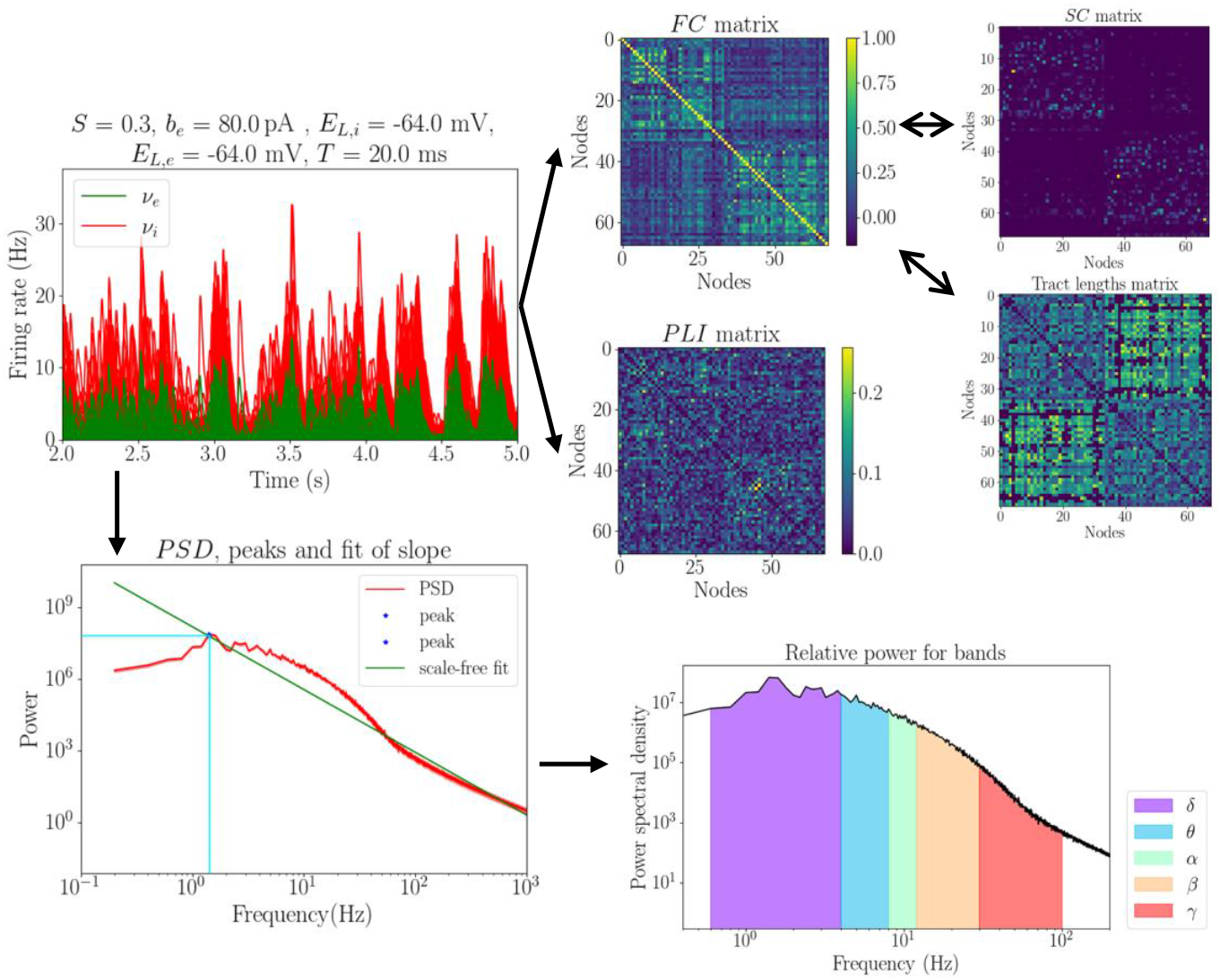
Feature Extraction pipeline. For each parameter combination, the features described in Tables 2 and 3 are obtained. All the features stem from the raw time-traces of the model, from which the FC matrix, the PLI matrix and the *PSD*(*f*) are obtained. The FC matrix is compared with both the SC matrix and the Tract Lengths matrix, and also used for a study on Default Mode Network correlations, explained below. Then, different algorithms are applied on the *PSD*(*f*) to obtain the spectral features.

**Figure 10:**
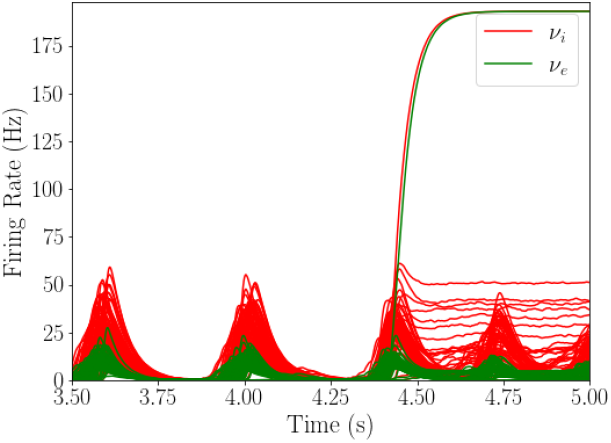
Example of how the *v*_*e*_ of one region in the TVB-AdEx model jumps to the pathological activity fixed point around 4.5 s.

**Figure 11:**
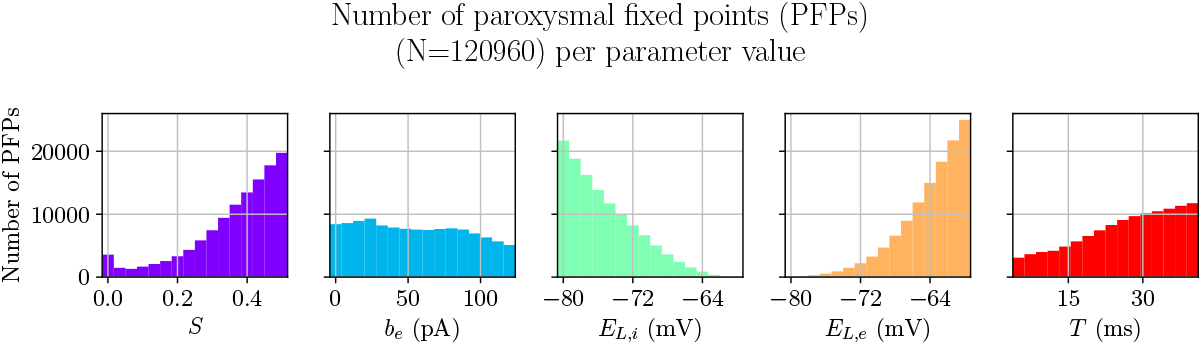
Histograms of number of times a Paroxysmal Fixed Point happens as a function of parameter values.

**Figure 12:**
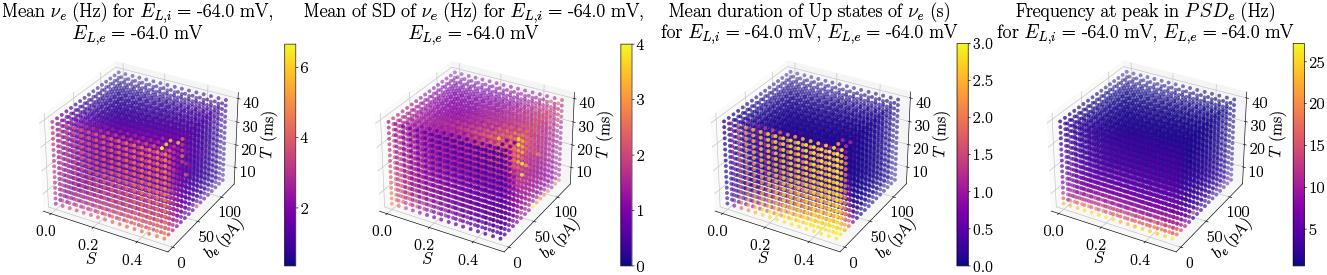
Features describing the behavior of the TVB-AdEx model in a subspace of the *depolarized region*, when varying *S, b*_*e*_ and *T*.

**Figure 13:**
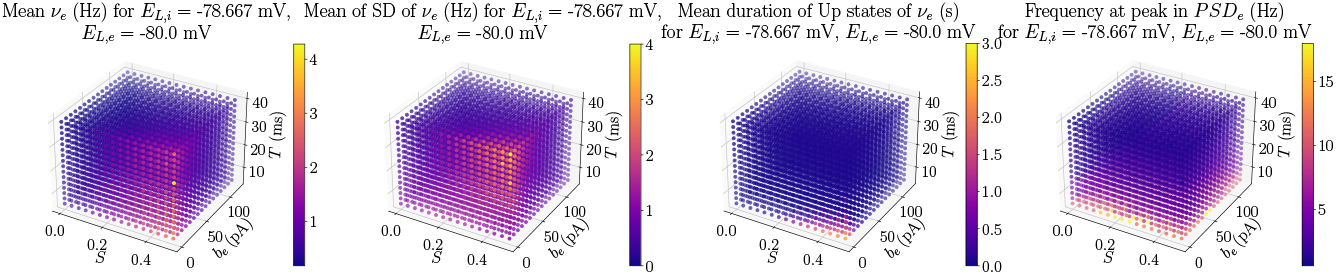
Features describing the behavior of the TVB-AdEx model in the *hyperpolarized region*, when varying *S, b*_*e*_ and *T*.

**Figure 14:**
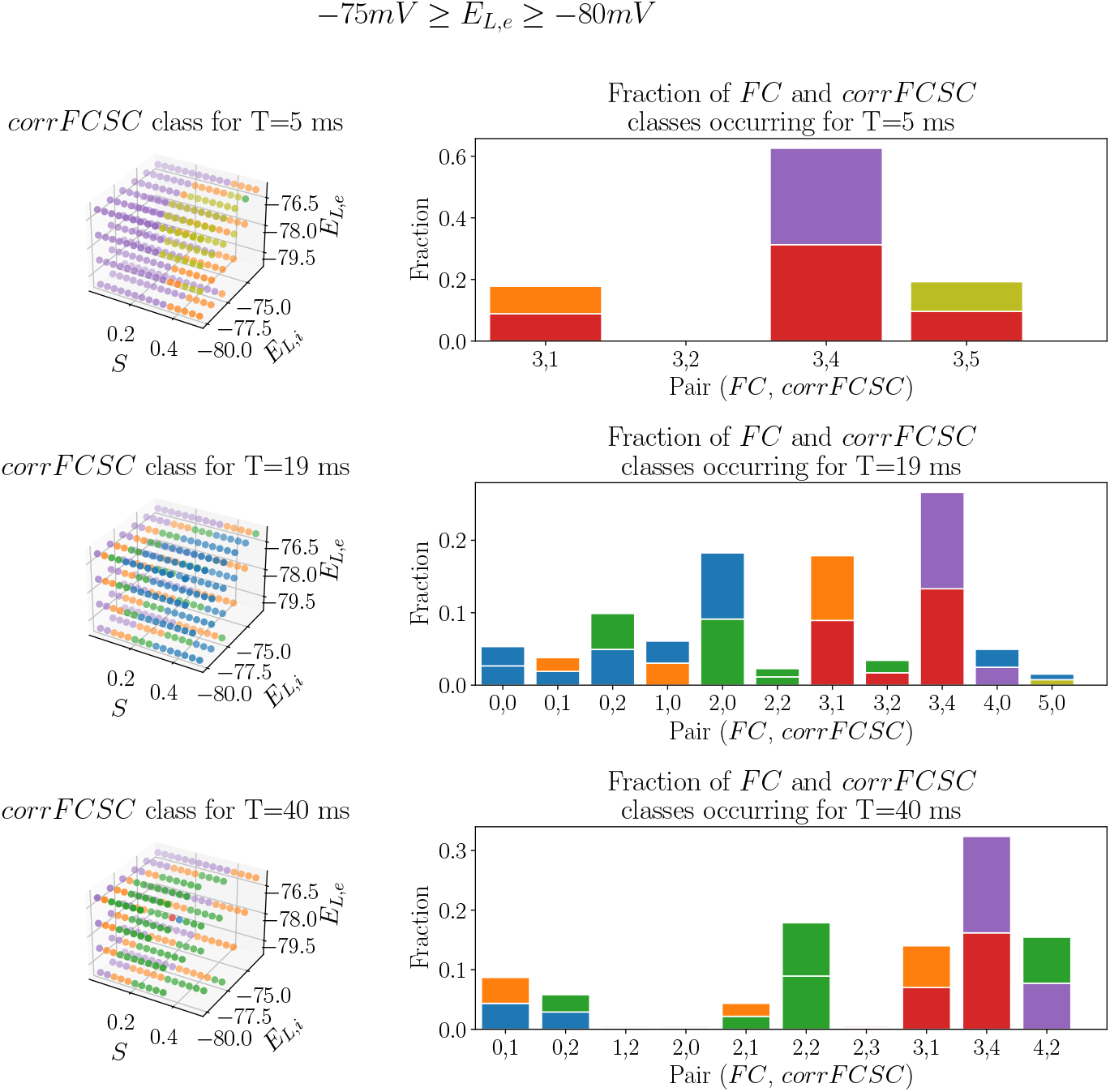
Distribution of *corrFCSC*(*b*_*e*_) classes as a function of *S, E*_*L,i*_ and *E*_*L,e*_ values for different *T* values (left panels) and histograms counting the fraction of times a point in the parameter space is assigned to *FC*(*b*_*e*_) class *i* and *corrFCSC*(*b*_*e*_) class *j* (right panels) for the *hyperpolarized* range of *E*_*L,e*_ (−75 mV ≥ *E*_*L,e*_ ≥ −80 mV). The bottom color of the histograms’ bars are the same as the pair’s *FC*(*b*_*e*_) color, while the top color is the same as the pair’s *corrFCSC*(*b*_*e*_) color.

#### A.2. Detailed Description of Features

### B. Supplementary results

#### B.1. Impact of parameter values on paroxysmal activity

To analyze the influence of each parameter on the appearance of Paroxysmal Fixed Points (PFPs) of the TVB-AdEx model, one can count, for all the parameter combinations containing a certain fixed parameter value, how many times the *v*_*e*_ threshold of 175 Hz is exceeded. That way, representations like the one in 11 can be built.

The excitatory transfer function of the AdEx mean field model (ℱ_*e*_ from Equations 1 and 2) is such that when it receives a high *v*_*e*_ input, the attractor can change, making the dynamics jump to the pathological fixed point at around 192 Hz. Thus, higher *v*_*e*_ incoming to the AdEx mean field models in the network will make it easier to reach this pathological activity. It is worth noting that a single AdEx mean-field model is an already complicated model to analyze dynamically. Propositions made in this appendix should be thoroughly tested through dynamical analyses in further work to better understand the mean-field model behavior.

Increased values of connectivity strength between nodes, *S*, result in higher *v*_*e*_ firing rates incoming to all the mean-field models in the network, considerably increasing the probability of reaching a PFP. Both leakage reversal potential, *E*_*L,i*_ and *E*_*L,e*_, play a similar but inverse role. When *E*_*L,i*_ takes lower values, the inhibitory populations of the network’s nodes become hyperpolarized, less active, leading to a higher excitatory activity, increasing the probability of going over the threshold for PFPs. When *E*_*L,e*_ takes lower values, the excitatory populations are less active, and the chances of reaching a PFP are lower, but when the leakage reversal potential increases, making excitatory populations more active, the probability rapidly increases. It is interesting to note how the *b*_*e*_ value has almost no influence on PFPs, since the probability of a PFP appearing remains almost constant for the whole *b*_*e*_ range. Finally, one can see in Figure 11 that PFP probability increases steadily with *T*. Increasing T makes the dynamical system slower in such a way that the trajectory of the *v*_*e*_ towards the fixed point is slower after a perturbation. Thus, incoming perturbations that move the trajectory towards higher activity can accumulate, each time increasing more and more the *v*_*e*_, until the transition line is crossed and the AdEx mean-field model is attracted by the 192 Hz fixed point.

#### B.2. Visualization of TVB-AdEx model’s evolution in time for certain parametrizations of the network

#### B.3. Relationship between FC and SC. Hyperpolarized Region

The behavior of the *hyperpolarized region* is considerably different to that of the *depolarized region*. In this case, it is hard to see homogeneous regions and the diversity of *FC*(*b*_*e*_), *corrFCSC*(*b*_*e*_) pairs is reduced. Again, for *T*= 5 ms, the only *FC*(*b*_*e*_) class available is the red one, indicating a total lack of overall connectivity in the TVB-AdEx network. In the *T* = 19 ms plots there is an interesting appearance of many blue and green *corrFCSC*(*b*_*e*_) points, which are related to those centroids that start at a considerably high level of *corrFCSC*. This is consistent with the fact that, in the *hyperpolarized region*, one can find UD states associated to unconscious dynamics even for low values of *b*_*e*_. Lastly, for *T* = 40 ms, the green *corrFCSC*(*b*_*e*_) class becomes the most frequent one again, as was the case of Figure 7, now being mostly paired with purple and green *FC*(*b*_*e*_) classes, which correspond to very high levels of *FC* throughout the *b*_*e*_ range.

## Notes

### Competing Interest Statement

The authors have declared no competing interest.

